# Genetic Information Processing Complexity as a Determinant of Virus Diversity

**DOI:** 10.64898/2026.06.01.729294

**Authors:** Shmuel Pietrokovski, Yosef Shaul

## Abstract

Viruses exhibit diverse genome architectures and replication strategies that shape their evolutionary trajectories and taxonomic diversification. Here, we test whether the complexity of viral genetic information processing predicts large-scale patterns of viral diversity. We define a propagation index that quantifies a minimal number of steps required for viral genome expression and replication across the Baltimore classes. Using ICTV taxonomy data (1971–2024), we identify a strong consistent linear relationship between the propagation index and viral diversification at both the family and genus levels. This statistically significant association is also observed for DNA and RNA viruses independently. Notably, the correlation persists across decades of ICTV releases despite substantial expansion and restructuring of viral taxonomy. Viruses with simpler propagation strategies consistently exhibit greater diversification, suggesting that genome processing complexity constrains macroevolutionary potential. These findings establish a quantitative link between propagation architecture and viral diversification and provide a predictive framework for understanding large-scale patterns of virus evolution.

## Introduction

Viruses are classified into taxonomic realms by their fundamental replication strategies (e.g., RNA vs. DNA genomes, reverse transcription). Further sub-classifications into kingdoms, phyla, and orders, is based on genome organization, replication strategy, and structural proteins (Black et al. 2025; Koonin et al. 2024, 2020). Orders often reflect adaptation to broad host categories (e.g., animals, plants, bacteria, archaea), and are further sub-classified into families, that are defined by genome size and structure, capsid shape, envelope presence, and replication mechanisms. Families are subdivided into genera, that contain closely related viruses, often infecting similar hosts or causing similar diseases. Genera include virus species that share high sequence similarity, host range, and pathogenicity.

Virus evolution involves their diversification and adaptation over time. A process shaped by mutation, recombination, natural selection, and host-virus interactions (Koonin and Dolja 2013; Sanjuán and Domingo-Calap 2016; Mifsud et al. 2025; Rasmussen and Stadler 2019). Unlike cellular organisms, viruses rely on the molecular machinery of their hosts, making their evolution highly dependent on mutation rates, recombination (Posada, Crandall, and Holmes 2002), reassortment, and selection pressures. These mechanisms drive viral diversification, influencing their evolutionary success and ability to persist in different environments.

Mutation is considered a primary driver of viral adaptation/evolution, particularly in RNA viruses, which exhibit high mutation rates due to error-prone replication by their RNA-dependent RNA polymerases (Dolja and Koonin 2018; Belshaw, Pybus, and Rambaut 2007; Sanjuán et al. 2010). In contrast, DNA viruses typically have more stable genomes but allow for alternative evolutionary strategies such as recombination. Genetic drift, the random accumulation of mutations, can produce neutral, beneficial, or deleterious effects on viral fitness (Duffy, Shackelton, and Holmes 2008).

Beyond mutation, genetic recombination and reassortment also contribute to viral diversity (Posada, Crandall, and Holmes 2002). Recombination occurs when two related viral genomes exchange genetic material, leading to novel variants, a process particularly common in DNA viruses and some RNA viruses. Reassortment, in contrast, happens in segmented RNA viruses (e.g., influenza), where genome segments are exchanged during co-infection, potentially giving rise to highly virulent or pandemic strains (Webster et al. 1982). These genetic processes, combined with selective pressures from host immune responses, antiviral treatments, and environmental factors, shape viral evolution. Positive selection favors mutations that enhance viral replication, transmission, or immune evasion, while negative selection eliminates deleterious mutations to maintain viral viability.

Viruses and their hosts exist in a continuous evolutionary arms race. Hosts develop immune defenses, while viruses evolve countermeasures, such as antigenic variation in influenza or immune evasion strategies in HIV (Karlsson Hedestam et al. 2008). This dynamic coevolution influences viral pathogenicity and persistence. Additionally, cross-species transmission plays a significant role in viral evolution, as seen in zoonotic viruses like SARS-CoV-2 and Ebola (Geoghegan, Duchêne, and Holmes 2017). These events are often facilitated by genetic changes that enhance host receptor binding and replication efficiency, leading to novel outbreaks and potential pandemics (Carabelli et al. 2023; Strobelt et al. 2023).

A growing body of data—especially over the last two decades—on the emergence and expansion of virus families presents an opportunity to evaluate viral evolution through a more quantitative lens. In light of clear patterns linking genome structure, replication strategies, and diversification, a fundamental question emerges: *Can aspects of viral evolution be quantitatively related to the diversity of different virus types?* Addressing this question may offer new insights into the predictability of viral emergence, the scope of yet unidentified viruses, and the broader dynamics of evolutionary biology. Here, we examine viral diversification at the family and genus levels and find they are quantitatively associated with the complexity of viral genome processing pathways.

## Methods and Rationale

### Virus classification and taxonomic data sources

Viruses were categorized according to their genome type and replication strategy using the Baltimore classification system, which groups viruses into seven classes based on the nature of their genetic material and the pathway they generate mRNA (Koonin et al. 2020; Baltimore 1971). These classes include double-stranded DNA (dsDNA), single-stranded DNA (ssDNA), double-stranded RNA (dsRNA), positive-sense single-stranded RNA (+ssRNA), negative-sense single-stranded RNA (−ssRNA), single-stranded RNA with reverse transcription (ssRNA-RT), and double-stranded DNA with reverse transcription (dsDNA-RT).

Taxonomic data were obtained from the International Committee on Taxonomy of Viruses (ICTV) Master Species Lists (MSLs), covering releases 1-40 from 1971 to 2024 (https://ictv.global/msl). For each release, we extracted the total number of recognized virus families and genera within each Baltimore class. We excluded from our analysis viroids, satellite viruses, virus-dependent nucleic acids, gene transfer agents, and any other that rely on helper viruses and/or are not categorized uniquely into one of the Baltimore classes (e.g., ambisense genome organization). This included 24/368 families and 82/3769 genera from the current 2024 release MSL 40v2 (supporting information tables s1-2).

Family and genera-level richness was used as the primary measure of macroevolutionary diversification, as these ranks capture deep evolutionary divergence while remaining relatively stable to frequent taxonomic reassignments at lower ranks (Black et al. 2025; Koonin et al. 2024, 2020). Genome compositions were taken directly from the ICTV MSL data, and prior to the MSL 29 release (1971-2013), by matching the families and genera with their corresponding taxon in later releases.

### Enumeration of genome processing steps

To quantify viral genetic processing complexity, we enumerated the minimal number of biochemical steps required for (i) viral protein expression and (ii) genome replication, for each Baltimore class. These steps were defined based on canonical replication pathways described in standard virology texts and reviews (Koonin et al. 2020).

A “processing step” was operationally defined as a major, obligate molecular transformation of nucleic acid required to produce functional viral proteins or progeny genomes. Examples include transcription of DNA into RNA, synthesis of complementary RNA strands, reverse transcription, and translation. Steps that are mechanistically similar but occur multiple times (e.g., repeated transcription events) were counted once, as the goal was to capture pathway complexity rather than kinetic frequency. Host-mediated processes that do not involve nucleic acid transformation (e.g., ribosome recruitment) were not counted as separate steps.

For each Baltimore class, the total number of translation-related and replication-related steps were determined independently (Fig. 1). These counts represent minimal processing pathways required for successful viral propagation under standard intracellular conditions.

**Figure 1.**
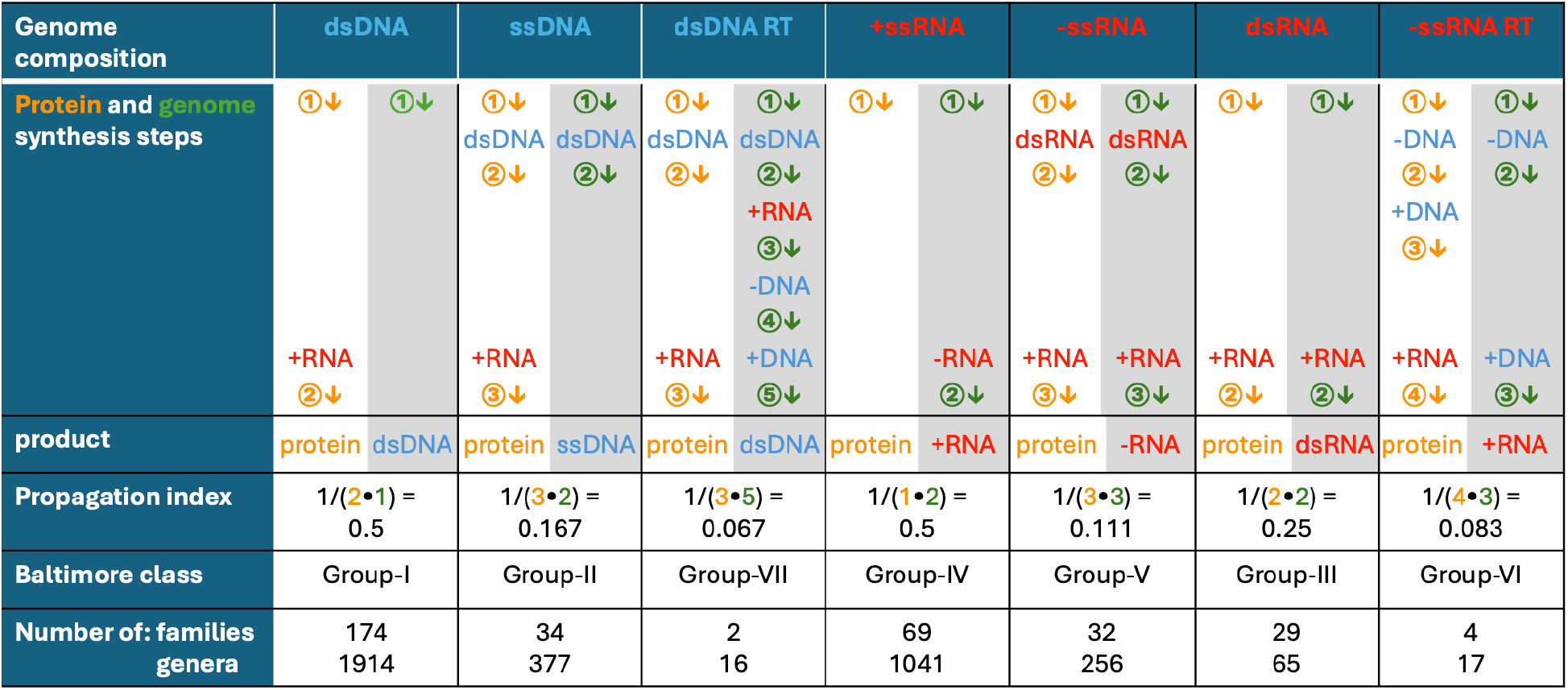
Viral genome processing complexity, propagation index, and taxonomic diversity across Baltimore viral classes. Schematic representation of the number of biochemical processing steps required for viral propagation, from the virus genome to protein production (orange) and genome synthesis (green, with gray background), across the different viral groups classified by the Baltimore system. Each numbered arrow indicates a distinct major biochemical step in replication or protein synthesis. The propagation index, calculated as the inverse of the product of replication and protein synthesis steps, quantifies the overall efficiency of genetic processing from replication to protein production. The number of analyzed viral families and genera is from the ICTV 2024 MSL 40v2 version.

### Definition of the propagation index

To integrate genome expression and replication complexity into a single quantitative metric, we defined a propagation index (PI) for each Baltimore class. The PI was calculated as the inverse of the product of the number of translation steps (T) and replication steps (R):

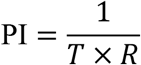

This formulation assigns higher PI values to viruses with simpler and more direct genetic processing pathways, and lower values to viruses requiring multiple intermediate conversions. The multiplicative structure reflects the assumption that expression and replication complexity jointly constrain propagation efficiency, rather than acting independently.

The propagation index is intended as a minimaly-based measure of genome processing complexity rather than a detailed kinetic or biochemical model. Each processing step is treated equivalently to capture the overall structure of the replication–expression pathway, independent of reaction rates, enzyme efficiencies, or host-specific factors. The step counts therefore provide a simplified, but consistent and tractable, framework for comparing virus reproduction complexity across Baltimore classes.

### Correlation analyses between propagation index and diversification

For each ICTV release, we examined the relationship between the propagation index and taxonomic diversification at the family and genus levels across Baltimore classes. Simple linear regression was used to assess the association between the PI and number of virus families or genera. Analyses were performed separately for DNA viruses, RNA viruses, and for all Baltimore classes combined, to account for known mechanistic differences between these groups.

For comparative analyses of the linear relations in multiple ICTV releases, each Baltimore class family and genus counts was normalized to its relative fraction, accounting for the overall growth in recognized virus diversity over time. This normalization ensured that correlations reflected relative diversification among classes rather than absolute increases driven by improved discovery and sequencing technologies. Coefficient of determination (R^2^) values were used for each regression to quantify the strength of the association.

### Permutation testing and statistical significance

To evaluate whether the observed correlations could arise by chance from the observed family or genus counts, we generated all possible 5040 (7!) permutations of the observed counts with the the propagation index values. Simple linear regression was calculated for each permutation, and the rank of the observed set in the R^2^ values gave an expected-value.

### Assumptions and limitations

This analysis assumes that Baltimore classes represent biologically meaningful units for comparing replication strategies and that amounts of family/genus-level are a reasonable proxy for macroevolutionary diversification (Koonin, Krupovic, and Agol 2021; Kitchen, Shackelton, and Holmes 2011; Koonin, Dolja, and Krupovic 2022). We note that taxonomic ranks are human-defined constructs that are influenced by sampling intensity, host bias, and evolving ICTV classification criteria. To evaluate this we examined the consistency of the observed relationships across decades of ICTV taxonomic revisions.

## Results

### Quantifying viral replication complexity reveals a predictive metric for diversification potential

Viral genome replication and gene expression vary significantly among the seven Baltimore classes (Koonin et al. 2020; Baltimore 1971), each requiring a different number of molecular steps to produce proteins (Fig. 1). These steps, which include reverse transcription, RNA synthesis, and translation, were systematically enumerated to capture the complexity of viral propagation. Steps needed to translate viral genomes into proteins, and the replication routes.“Simpler” classes such as positive-sense single-stranded RNA (+RNA) viruses translate directly into proteins with minimal processing, whereas retroviruses and reverse-transcribing DNA viruses require multiple enzymatic conversions and intermediates.

To quantify this complexity, we defined a *propagation index* as the inverse product of the number of replication and translation steps (Fig. 1). This metric assigns higher values to simpler viruses and lower values to more complex ones. When compared to the ICTV 2022 taxonomy data (Fig. 1), a clear trend emerges: virus classes with higher propagation indices tend to include a greater number of families. For instance, +RNA and dsDNA viruses, which require fewer processing steps, exhibit the largest family and genus counts. In contrast, retroRNA and retroDNA viruses, with more elaborate replication routes, display the lowest propagation indices and the smallest number of families and genera.

This analysis suggests that viral propagation simplicity may provide an evolutionary advantage by enhancing replication efficiency and resilience against host defenses, allowing broader diversification. The propagation index can thus serve as a quantitative tool to predict viral diversification potential across virus types.

### The number of virus families is linearly correlated with the propagation index across DNA and RNA viruses (ICTV 2024 data)

To test whether propagation index predicts viral diversification, we examined its relationship with the number of virus families using ICTV classification data. The latest 2024 data DNA (Fig. 2A) and RNA viruses (Fig. 2B) each show strong linear correlations (R^2^: 0.998 and 0.837, respectively). A clear but lower correlation (R^2^: 0.699) is present when analyzing all Baltimore viral classes (Fig. 2C). This correlation supports our hypothesis, and its lower value from that of the DNA and RNA viruses each potentially reflects mechanistic differences between DNA and RNA viruses or differences in sampling and classification.

**Figure 2.**
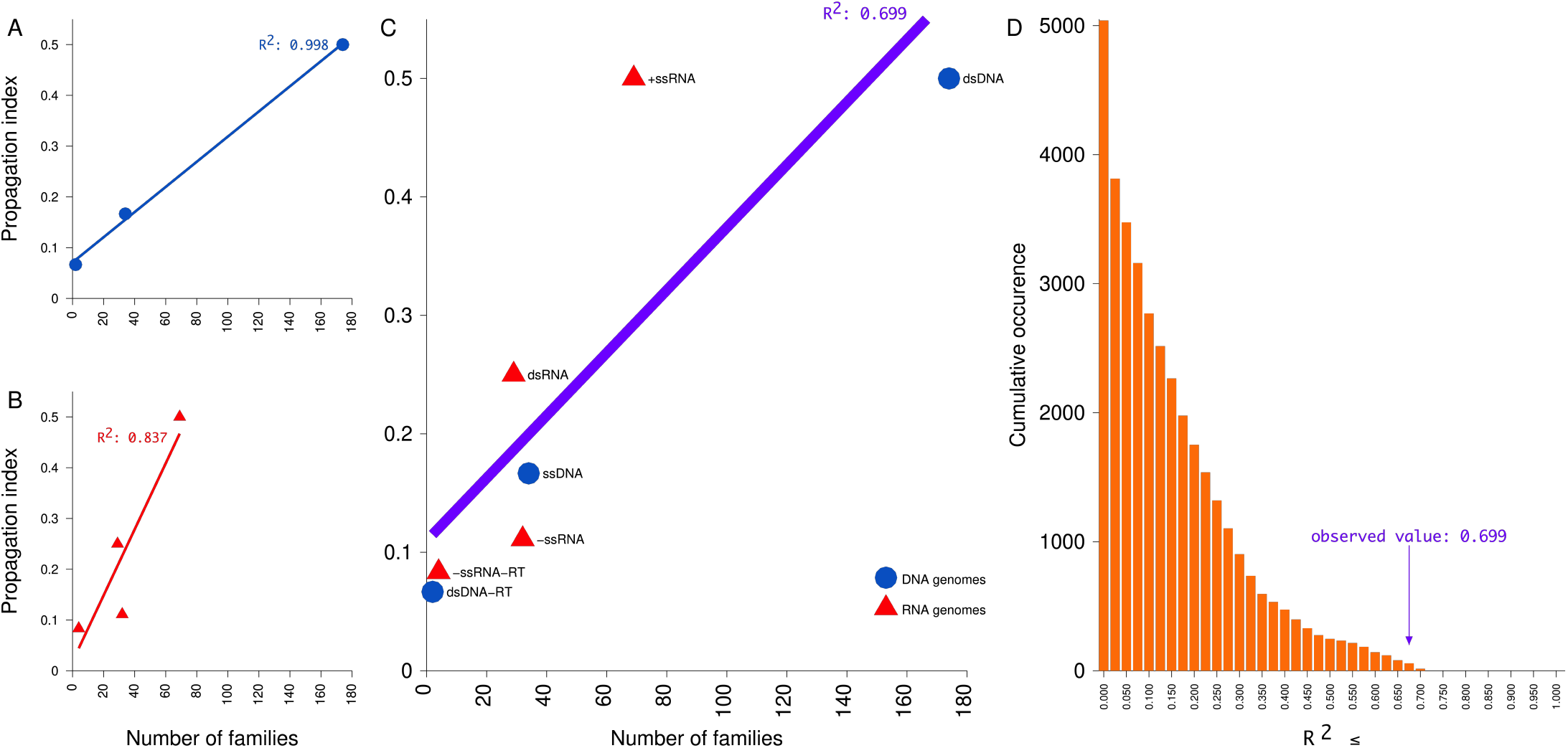
Relationship between propagation index and the number of virus families. (A–C) Scatter plots showing the linear relationship between propagation index (y-axis) and number of virus families (x-axis) for (A) DNA viruses, (B) RNA viruses, and (C) DNA and RNA viruses, from all Baltimore classes. Each point represents a virus class, with DNA viruses in blue and RNA viruses in red. The linear regression line is shown for each set together with its R^2^ value. (D) Cumulative occurrence histogram of R^2^ values from all possible combinations of the family counts and propagation index of the data shown in panel C. Each bar shows how many values are less or equal to an R^2^ value. The actual observed R^2^ value is marked. It is the 18/5040 highest value. The histogram of the raw values used to create this cummulative histogram is in figure s1A.

To assess the significance of this correlation, we calculated the same analysis for all possible permutations of the family counts and propagation index values (Fig. 2D). A total of 5,040 permutations were performed, generating a null distribution of R^2^ values under the assumption of no relationship between genetic processing complexity and diversification. The empirical R^2^ obtained from the observed data was compared to this distribution to estimate statistical significance. Observed R^2^ values lying in the extreme tail of the null distribution were interpreted as evidence against the null hypothesis of random association. Only 18/5040 of these permuted sets had higher or identical R^2^ values than the observed one, giving an expected value of 3.57•10^−3^. This indicates that the correlation between propagation index and family diversity is unlikely to have occurred by chance from the observed values. These findings support the hypothesis that simpler propagation strategies are associated with greater viral diversification, possibly by evading multiple host checkpoints.

### Propagation index predicts viral genus-level diversification

To determine whether the above relationship is present in a finer taxonomic resolution, we examined the association between propagation index and viral genus diversity in the same way. As above, DNA viruses exhibited an almost perfect linear relationship between the number of genera and the propagation index (R^2^ = 0.998; Figure 3A) and RNA viruses showed a strong association, although with greater dispersion (R^2^ = 0.787; Figure 3B). Analysis of all the viruses together also gave a strong correlation (R^2^ = 0.775; Figure 3C), indicating that the relationship is robust and is not restricted to specific virus domains.

**Figure 3.**
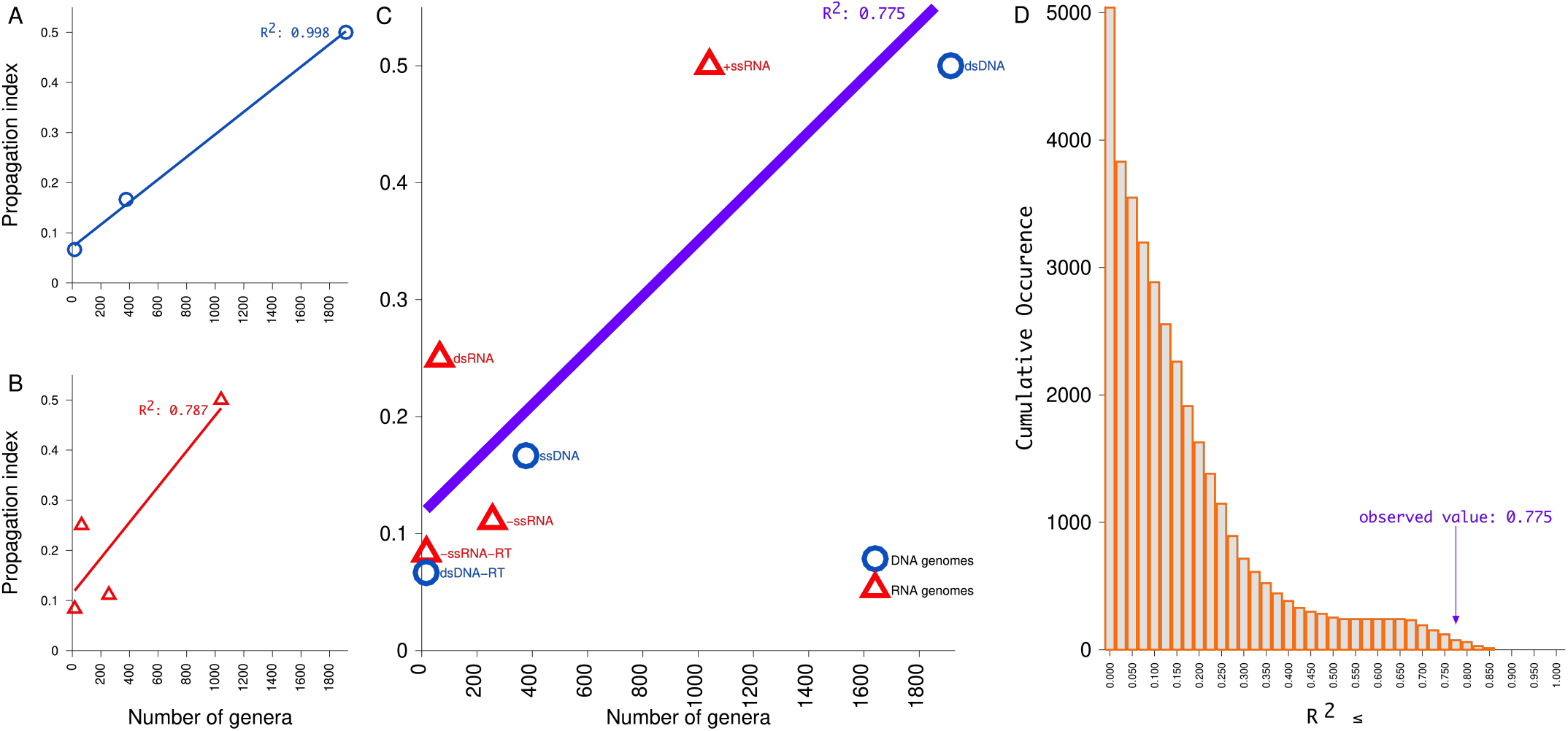
Relationship between propagation index and the number of virus genera. Analyses and data for ICTV genera are shown as in figure 2.

As above, to assess whether this correlation could arise by chance, we examined all possible permutation of the propagation index values with the counts of the viral groups. The observed R^2^ value of 0.775 fell in the extreme upper tail of the resulting null distribution and gave an e-value of 1.47•10^−2^ (74/5040; Figure 3D), supporting the statistical significance and robustness of the association. Together, these results indicate that the link between propagation index and viral diversification at the genus level is strong and unlikely to reflect random variation.

### A consistent relationship between propagation index and virus family and genera diversification persists over more then three decades

The total number of recognized virus families and genera has increased steadily from 1971 to 2024, reflecting ongoing viral discovery and refinement of classification criteria by the ICTV (Fig 4A and D). To determine whether the correlation between viral replication complexity and taxonomic diversification holds over time, we analyzed all ICTV releases from version 12 in 1991 to the latest version 40 in 2024. Previous releases have lower amounts of families, genera, and other taxons and are thus more prone to sampling fluctuations. The linear correlations between the propagation index and the number of virus families and genera across all Baltimore classes in each release have similar slopes (Fig 4B and E), and all have consistently high R^2^ values (Fig 4C and F). Thus, the linear relationship we describe is stable for both families and genera across decades of ICTV data and classifications, despite substantial taxonomic growth and restructuring. These results support the hypothesis that simpler propagation mechanisms enable broader viral diversification and that this principle is robust to historical changes in virus classification.

**Figure 4.**
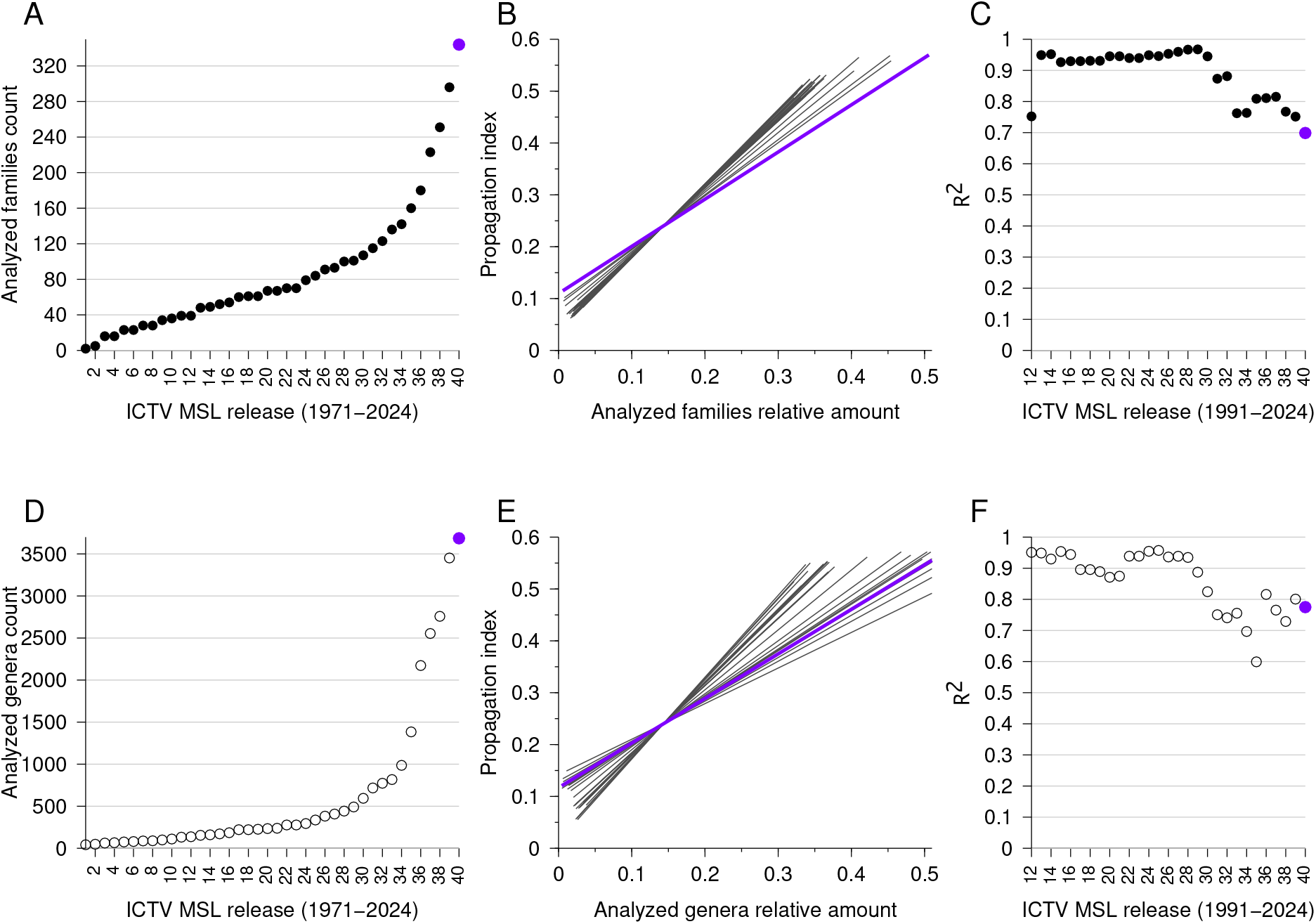
Long-term consistency in the correlation between propagation index and virus family and genus diversity across ICTV releases (1971–2024). (A) The number of analyzed virus families (e.g., excluding viroids and satellite viruses, see section ‘Virus classification and taxonomic data sources’) defined in each ICTV Master Species List (MSL) release. (B) Linear regression fits between the propagation index and normalized number (relative amount) of families for each MSL release from 1991 (when the total number of families was 39) to 2024. The 2024 release (MSL 40) is highlighted in purple. (C) R^2^ values for each regression line in B. To compare the lines derived from different MSL releases the taxon (families or genera) counts were normalized to the fractions of the total number of the taxon, giving relative amounts. This does not change the line intercept, R or R^2^ measures, only the slope. (D-F) data and analyses of genera as in A-C.

## Discussion

Our analysis identifies a robust and consistent relationship between viral genetic processing complexity and taxonomic diversification at the family and genus level. By quantifying the number of molecular steps required for genome replication and gene expression, we introduce a propagation index that captures a fundamental aspect of viral biology. Across all Baltimore classes, viruses with simpler propagation strategies consistently exhibit greater diversification, a significant pattern that persists across decades of ICTV taxonomic revisions. This consistency strongly suggests that the observed correlation reflects an underlying evolutionary constraint rather than a transient artifact of virus discovery or classification practices. Finding the same results on both family and genus taxonomic levels strengthens our findings. The number of virus genera is not trivialy derived from the number of families. While typically most families in the latest ICTV release include 1-2 genera, many have up to 70 genera and 3 +ssRNA families have 123, 202, and 308 genera (Atkinsviridae, Steitzviridae, and Fiersviridae, respectively; Supporting information figure S2).

The propagation index represents a simplified abstraction of viral replication complexity and does not explicitly account for kinetic parameters, error rates, or host-specific factors.Nevertheless, the persistence of the observed correlation across multiple ICTV releases and alternative formulations suggests that the relationship reflects a general constraint associated with genome processing architecture rather than a modeling artifact or bias.

The relationship is particularly strong among DNA viruses, where replication fidelity and genome stability are relatively high. RNA viruses, in contrast, display greater variability around the regression, likely reflecting the combined effects of high mutation rates, error-prone replication, and frequent population bottlenecks (Duffy, Shackelton, and Holmes 2008; Sanjuán and Domingo-Calap 2016). While elevated mutation rates promote rapid adaptation and short-term evolutionary success, they may also impose limits on long-term lineage stability through mutational load and extinction risk (Regoes, Hamblin, and Tanaka 2013; Peck and Lauring 2018). This tradeoff may partially explain why replication simplicity alone is a less precise predictor of diversification among RNA viruses.

Several alternative explanations must also be considered. One possibility is uneven sampling and historical bias in virus discovery and classification (Edwards and Rohwer 2005). RNA viruses— particularly those infecting humans, livestock, and crops—have been disproportionately studied due to their medical and economic importance, while many DNA viruses infecting understudied hosts or environmental reservoirs remain poorly characterized (Simmonds et al. 2017). Advances in metagenomics have further reshaped virus taxonomy, often expanding family and genus levels diversity unevenly across Baltimore classes (Simmonds et al. 2017; Black et al. 2025).Nevertheless, the persistence of the correlation across multiple ICTV releases—despite major shifts in sequencing technologies, sampling strategies, and taxonomic organization criteria— argues against sampling bias as the sole explanation.

Ecological factors may further modulate viral diversification. Host range, transmission mode, and opportunities for cross-species transmission all influence evolutionary trajectories (Koonin and Dolja 2013; Koonin et al. 2024). Viruses with simpler propagation strategies may more readily adapt to novel hosts, increasing ecological breadth and opportunities for diversification. In this context, propagation simplicity may not act in isolation but instead synergize with ecological flexibility to expand the accessible evolutionary landscape. Host–virus coevolution, immune pressure, and cross-species transmission are therefore likely to interact with replication architecture to shape long-term macroevolutionary patterns (Longdon et al. 2014; Kaján et al. 2020; Hulse et al. 2023).

Our use of family and genera-level richness as a proxy for evolutionary success also warrants consideration. Taxonomic ranks are human-defined constructs that reflect both biological reality and historical convention (Caetano-Anollés, Claverie, and Nasir 2023). While family and genera counts provide a broadly comparable measure of macroevolutionary diversification, alternative metrics—such as lineage age, total species richness, or phylogenetic diversity—may capture complementary dimensions of evolutionary success. Future studies integrating the propagation index with time-calibrated phylogenies and alternative diversity measures will further test the generality and predictive power of the proposed framework.

Despite these limitations, the propagation index offers a simple and quantitative link between genomic information proccessing and large-scale evolutionary outcomes. Rather than implying a deterministic rule, our findings support a constraint-based model in which genetic processing complexity shapes the probability space of viral evolution. Simpler propagation strategies appear to relax constraints on diversification, increasing the likelihood of lineage expansion over macroevolutionary timescales. This view aligns viral macroevolution with broader evolutionary principles, in which reduced molecular or developmental complexity is often associated with increased evolvability (Strobel, Horwitz, and Meyer 2022; Strobel, Stuart, and Meyer 2022; Payne and Wagner 2019).

In summary, our analyses suggest that viral macroevolution is shaped not only by mutation rates, host interactions, and ecological opportunity, but also by the fundamental architecture of genome processing itself. By quantitatively linking replication strategy to taxonomic diversification, this study provides a unifying framework for interpreting viral evolutionary potential and offers a predictive perspective on the long-term consequences of viral genome organization.

## Supplementary information

**Figure S1.**
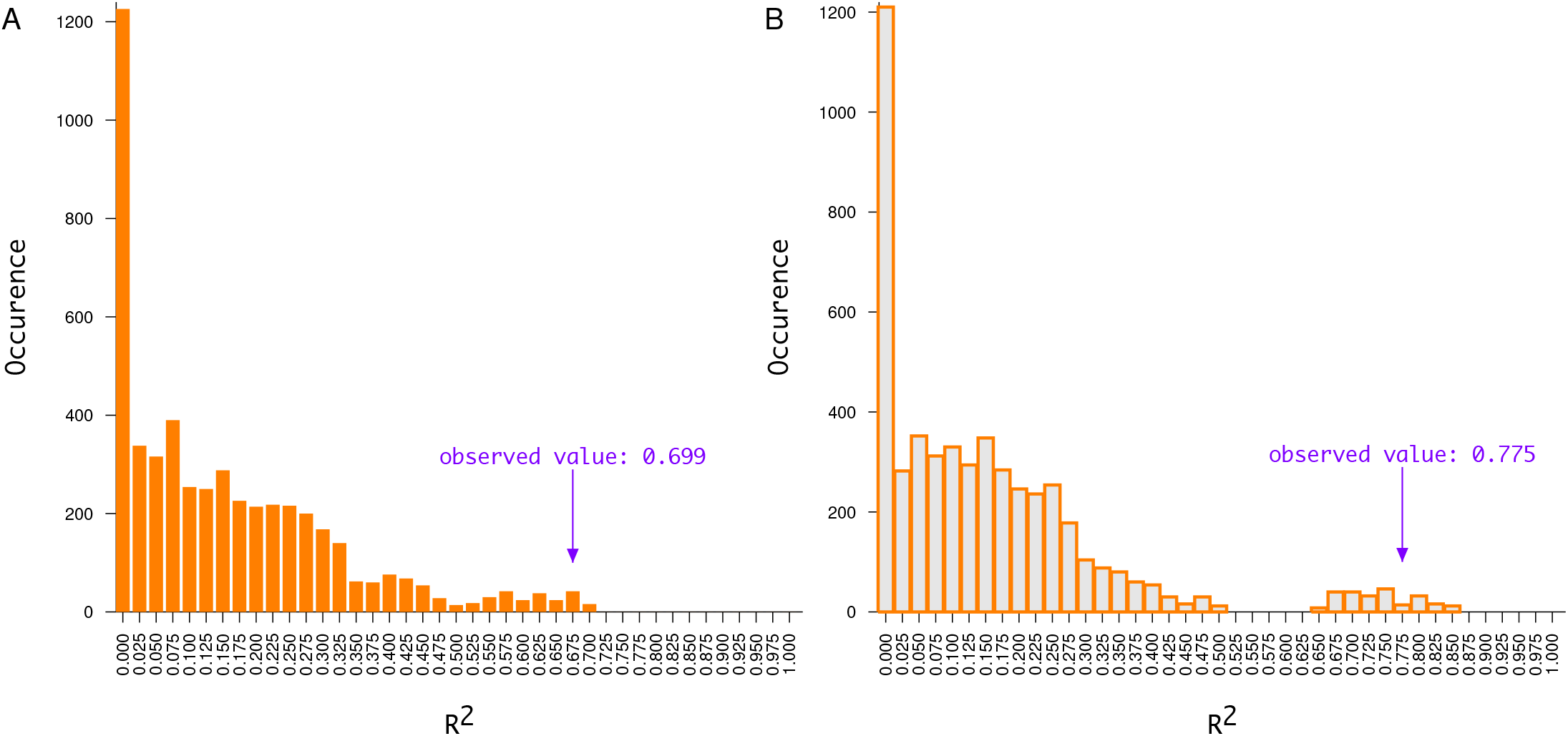
Occurrence histograms of R^2^ values from all possible combinations of family and genera counts with propagation index values. Histogram of R^2^ values from all possible combinations of the family (A) and genera (B) counts and propagation index of the data shown in panels C of figures 2 and 3. The actual observed R^2^ values are marked. The cumulative histograms of these values are in panels D of figures 2 and 3.

**Figure S2.**
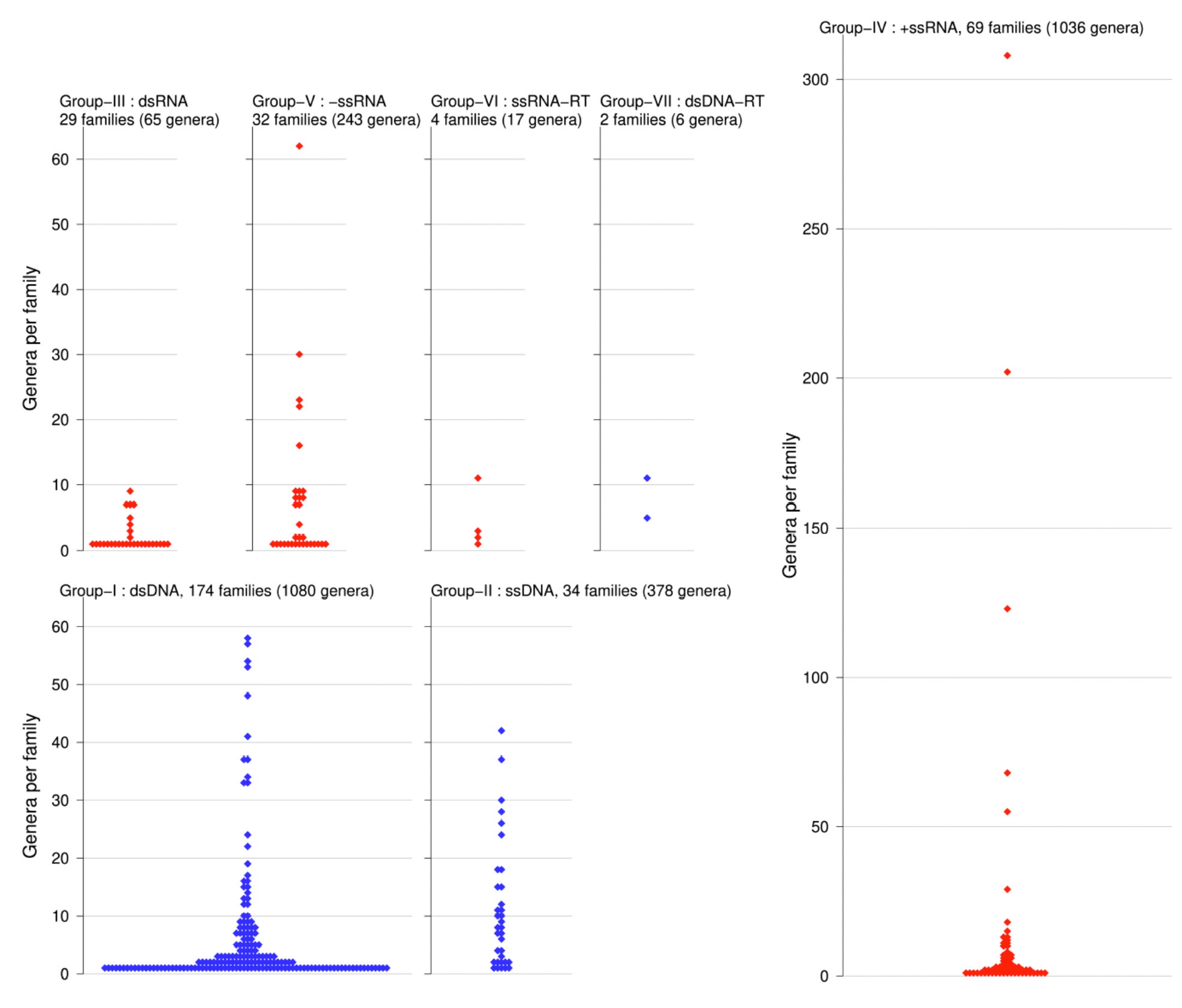
Number of genera per analyzed family across each Baltimore class. Beeswarm histograms, where each mark is a family with the number genera shown in the Y-axis. The total number of families and genera is shown for each Baltimore class.

**Table S1.**
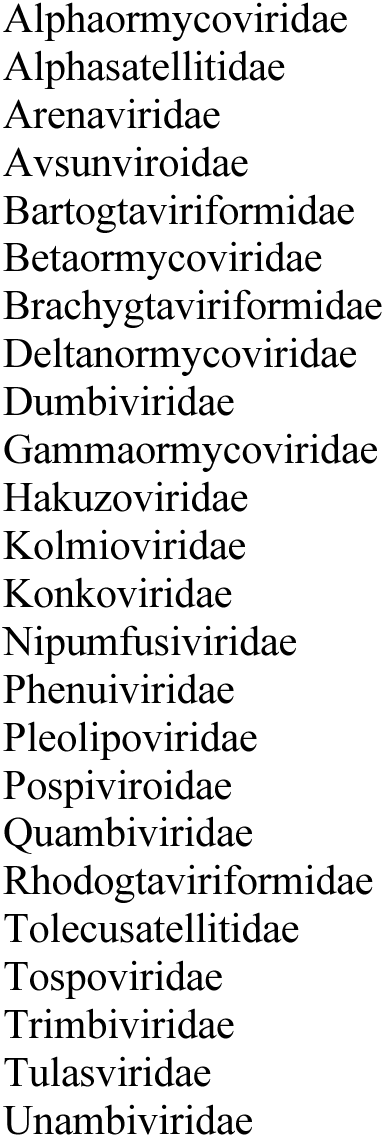
excluded ICTV families.

**Table S2.**
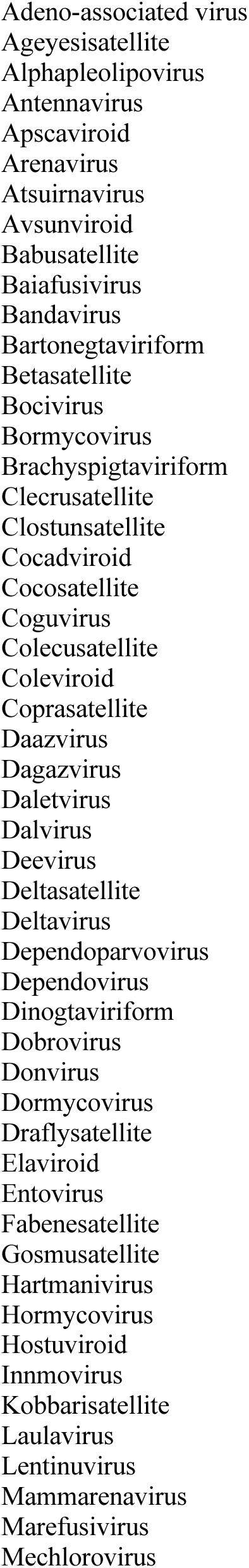
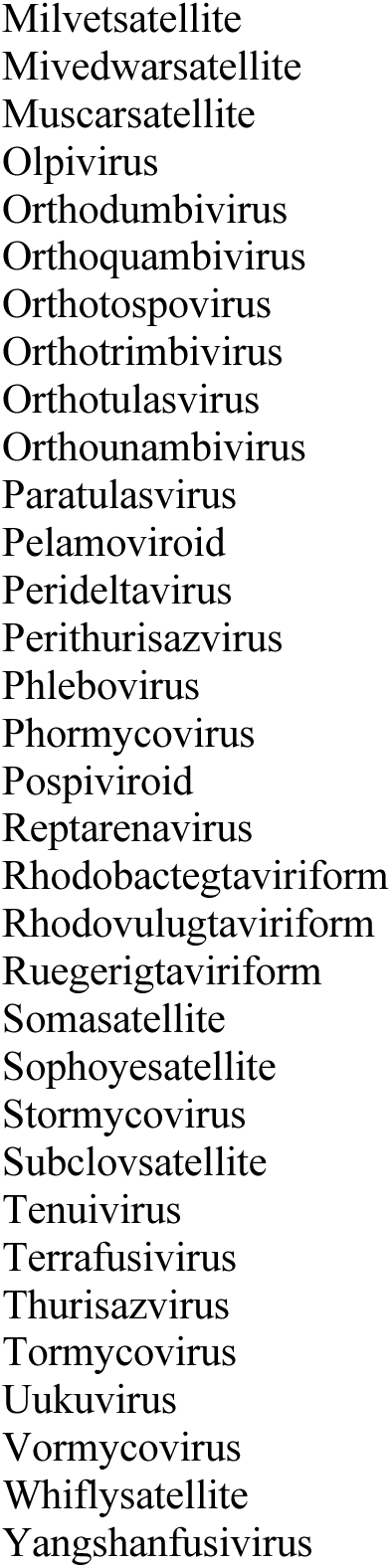
excluded ICTV genera.

## References

Baltimore, D. 1971. “Expression of Animal Virus Genomes.” Bacteriological Reviews 35 (3): 235–41. 10.1128/br.35.3.235-241.1971.

Belshaw, Robert, Oliver G Pybus, and Andrew Rambaut. 2007. “The Evolution of Genome Compression and Genomic Novelty in RNA Viruses.” Genome Research 17 (10): 1496–1504. 10.1101/gr.6305707.

Black, Eden J, C Steve Powell, Donald M Dempsey, R Curtis Hendrickson, Logan R Mims, and Elliot J Lefkowitz. 2025. “Virus Taxonomy: The Database of the International Committee on Taxonomy of Viruses.” Nucleic Acids Research, November. 10.1093/nar/gkaf1159.

Caetano-Anollés, Gustavo, Jean-Michel Claverie, and Arshan Nasir. 2023. “A Critical Analysis of the Current State of Virus Taxonomy.” Frontiers in Microbiology 14 (August): 1240993. 10.3389/fmicb.2023.1240993.

Carabelli, Alessandro M, Thomas P Peacock, Lucy G Thorne, William T Harvey, Joseph Hughes, COVID-19 Genomics UK Consortium, Sharon J Peacock, et al. 2023. “SARS-CoV-2 Variant Biology: Immune Escape, Transmission and Fitness.” Nature Reviews Microbiology 21 (3): 162–77. 10.1038/s41579-022-00841-7.

Dolja, Valerian V, and Eugene V Koonin. 2018. “Metagenomics Reshapes the Concepts of RNA Virus Evolution by Revealing Extensive Horizontal Virus Transfer.” Virus Research 244 (January): 36–52. 10.1016/j.virusres.2017.10.020.

Duffy, Siobain, Laura A Shackelton, and Edward C Holmes. 2008. “Rates of Evolutionary Change in Viruses: Patterns and Determinants.” Nature Reviews. Genetics 9 (4): 267–76. 10.1038/nrg2323.

Edwards, Robert A, and Forest Rohwer. 2005. “Viral Metagenomics.” Nature Reviews Microbiology 3 (6): 504–10. 10.1038/nrmicro1163.

Geoghegan, Jemma L, Sebastián Duchêne, and Edward C Holmes. 2017. “Comparative Analysis Estimates the Relative Frequencies of Co-Divergence and Cross-Species Transmission within Viral Families.” PLoS Pathogens 13 (2): e1006215. 10.1371/journal.ppat.1006215.

Hulse, Samuel V, Janis Antonovics, Michael E Hood, and Emily L Bruns. 2023. “Host-Pathogen Coevolution Promotes the Evolution of General, Broad-Spectrum Resistance and Reduces Foreign Pathogen Spillover Risk.” Evolution Letters 7 (6): 467–77. 10.1093/evlett/qrad051.

Kaján, Győző L, Andor Doszpoly, Zoltán László Tarján, Márton Z Vidovszky, and Tibor Papp. 2020. “Virus-Host Coevolution with a Focus on Animal and Human DNA Viruses.” Journal of Molecular Evolution 88 (1): 41–56. 10.1007/s00239-019-09913-4.

Karlsson Hedestam, Gunilla B, Ron A M Fouchier, Sanjay Phogat, Dennis R Burton, Joseph Sodroski, and Richard T Wyatt. 2008. “The Challenges of Eliciting Neutralizing Antibodies to HIV-1 and to Influenza Virus.” Nature Reviews Microbiology 6 (2): 143–55. 10.1038/nrmicro1819.

Kitchen, Andrew, Laura A Shackelton, and Edward C Holmes. 2011. “Family Level Phylogenies Reveal Modes of Macroevolution in RNA Viruses.” Proceedings of the National Academy of Sciences of the United States of America 108 (1): 238–43. 10.1073/pnas.1011090108.

Koonin, Eugene V, Valerian V Dolja, Mart Krupovic, Arvind Varsani, Yuri I Wolf, Natalya Yutin, F Murilo Zerbini, and Jens H Kuhn. 2020. “Global Organization and Proposed Megataxonomy of the Virus World.” Microbiology and Molecular Biology Reviews 84 (2). 10.1128/MMBR.00061-19.

Koonin, Eugene V, Valerian V Dolja, and Mart Krupovic. 2022. “The Logic of Virus Evolution.” Cell Host & Microbe 30 (7): 917–29. 10.1016/j.chom.2022.06.008.

Koonin, Eugene V, and Valerian V Dolja. 2013. “A Virocentric Perspective on the Evolution of Life.” Current Opinion in Virology 3 (5): 546–57. 10.1016/j.coviro.2013.06.008.

Koonin, Eugene V, Mart Krupovic, and Vadim I Agol. 2021. “The Baltimore Classification of Viruses 50 Years Later: How Does It Stand in the Light of Virus Evolution?” Microbiology and Molecular Biology Reviews 85 (3): e0005321. 10.1128/MMBR.00053-21.

Koonin, Eugene V, Jens H Kuhn, Valerian V Dolja, and Mart Krupovic. 2024. “Megataxonomy and Global Ecology of the Virosphere.” The ISME Journal 18 (1). 10.1093/ismejo/wrad042.

Longdon, Ben, Michael A Brockhurst, Colin A Russell, John J Welch, and Francis M Jiggins. 2014. “The Evolution and Genetics of Virus Host Shifts.” PLoS Pathogens 10 (11): e1004395. 10.1371/journal.ppat.1004395.

Mifsud, Jonathon C O, Marc A Suchard, Edward C Holmes, and Philippe Lemey. 2025. “Recent Advances in the Inference of Deep Viral Evolutionary History.” Journal of Virology 99 (9): e0029225. 10.1128/jvi.00292-25.

Payne, Joshua L, and Andreas Wagner. 2019. “The Causes of Evolvability and Their Evolution.” Nature Reviews. Genetics 20 (1): 24–38. 10.1038/s41576-018-0069-z.

Peck, Kayla M, and Adam S Lauring. 2018. “Complexities of Viral Mutation Rates.” Journal of Virology 92 (14). 10.1128/JVI.01031-17.

Posada, David, Keith A Crandall, and Edward C Holmes. 2002. “Recombination in Evolutionary Genomics.” Annual Review of Genetics 36 (June): 75–97. 10.1146/annurev.genet.36.040202.111115.

Rasmussen, David A, and Tanja Stadler. 2019. “Coupling Adaptive Molecular Evolution to Phylodynamics Using Fitness-Dependent Birth-Death Models.” ELife 8 (August). 10.7554/eLife.45562.

Regoes, Roland R, Steven Hamblin, and Mark M Tanaka. 2013. “Viral Mutation Rates: Modelling the Roles of within-Host Viral Dynamics and the Trade-off between Replication Fidelity and Speed.” Proceedings. Biological Sciences / the Royal Society 280 (1750): 20122047. 10.1098/rspb.2012.2047.

Sanjuán, Rafael, and Pilar Domingo-Calap. 2016. “Mechanisms of Viral Mutation.” Cellular and Molecular Life Sciences 73 (23): 4433–48. 10.1007/s00018-016-2299-6.

Sanjuán, Rafael, Miguel R Nebot, Nicola Chirico, Louis M Mansky, and Robert Belshaw. 2010. “Viral Mutation Rates.” Journal of Virology 84 (19): 9733–48. 10.1128/JVI.00694-10.

Simmonds, Peter, Mike J Adams, Mária Benkő, Mya Breitbart, J Rodney Brister, Eric B Carstens, Andrew J Davison, et al. 2017. “Consensus Statement: Virus Taxonomy in the Age of Metagenomics.” Nature Reviews Microbiology 15 (3): 161–68. 10.1038/nrmicro.2016.177.

Strobelt, Romano, Karin Broennimann, Julia Adler, and Yosef Shaul. 2023. “SARS-CoV-2 Omicron Specific Mutations Affecting Infectivity, Fusogenicity, and Partial TMPRSS2-Independency.” Viruses 15 (5). 10.3390/v15051129.

Strobel, Hannah M, Elijah K Horwitz, and Justin R Meyer. 2022. “Viral Protein Instability Enhances Host-Range Evolvability.” PLoS Genetics 18 (2): e1010030. 10.1371/journal.pgen.1010030.

Strobel, Hannah M, Elizabeth C Stuart, and Justin R Meyer. 2022. “A Trait-Based Approach to Predicting Viral Host-Range Evolvability.” Annual Review of Virology 9 (1): 139–56. 10.1146/annurev-virology-091919-092003.

Webster, R G, W G Laver, G M Air, and G C Schild. 1982. “Molecular Mechanisms of Variation in Influenza Viruses.” Nature 296 (5853): 115–21. 10.1038/296115a0.

